# Deciphering the genetic and epidemiological landscape of mitochondrial DNA abundance

**DOI:** 10.1101/2020.09.25.313171

**Authors:** Sara Hägg, Juulia Jylhävä, Yunzhang Wang, Kamila Czene, Felix Grassmann

## Abstract

Somatically acquired whole chromosome loss in nucleated blood cells is an indicator of immune aging and genomic instability. However, little is known about aging, lifestyle and genetic factors influencing mitochondrial (MT) DNA abundance. In this study, MT DNA abundance was estimated from the weighted intensities of probes mapping to the MT genome in 295,150 participants from the UK Biobank. We found that abundance of MT DNA was significantly elevated in women compared to men, was negatively correlated with advanced age, more packyears, greater body-mass index, higher frailty index as well as elevated red and white blood cell count and, importantly, lower mortality. In addition, several biochemistry markers in blood related to cholesterol metabolism, ion homeostasis and kidney function were found to be significantly associated with MT DNA abundance. By performing a genome-wide association study, we identified 50 independent regions genome-wide significantly associated with MT DNA abundance which harbour multiple genes involved in the immune system, cancer as well as mitochondrial function. Using mixed effects models, we estimated the SNP-heritability of MT DNA abundance to be around 8%. To investigate the consequence of altered MT DNA abundance, we performed a phenome-wide association study and found that MT DNA abundance is involved in risk for leukaemia, hematologic diseases as well as hypertension. Thus, estimating MT DNA abundance from genotyping arrays has the potential to provide novel insights into age- and disease relevant processes, particularly those related to immunity and established mitochondrial functions.

## Introduction

Mitochondria (MT), in addition to producing energy through oxidative processes, are involved in heat production, ion storage, apoptosis, intra- and extra-cellular cell signalling, biosynthesis and degradation of important metabolites as well as processing of therapeutic agents. Mitochondria have their own protein apparatus encoded in a small genome (16,569 base pairs). In contemporary human populations, several haplogroups are defined by ancestral and stable mutations in the MT genome. Those haplogroups are derived from adaptation to different geographic areas under distinctive selection pressure ^1^. Notably, different haplogroups have been reported to be associated with diseases as well as longevity ^2, 3^ and as such contribute to phenotypic variety within human populations. Mitochondria rely on self-replication and are thus prone to cellular stress in aging and diseases. Hence, MT dysfunction is a hallmark of aging ^4^, and has been associated with most aging-related diseases ^5^ as well as immunological processes^6^.

Another hallmark of aging is genomic instability, which can lead to loss of genetic material frequently observed in most cell types in the aging body ^4^. The best studied and most common type of somatic (whole) chromosomal loss in humans is the mosaic loss of Y chromosome in men, which has been correlated to increased risk of numerous non-haematological diseases as well as all-cause mortality ^7–11^. The percentage of nucleated cells in blood that do not carry a Y chromosome defines the degree of Y chromosome loss, which can vary strongly across male populations. Recent studies have analysed the intensities from hundreds or thousands of probes on high-resolution genotyping arrays to measure the abundance of sex chromosomes in circulating blood cells and found that Y chromosome loss is mainly influenced by advanced age, smoking and higher body mass index (BMI) ^11–13^. In contrast to Y chromosome loss, we have limited knowledge whether the abundance of other chromosomes changes with advanced age or due to environmental or life-style factors. However, investigating the underpinnings of chromosomal loss or gain is paramount to understand and potentially treat processes involved in aging and diseases.

To our knowledge, there is presently no comprehensive evaluation of aging or other factors influencing MT DNA abundance in nucleated blood cells. In addition, few studies have been conducted to elucidate the inherited genetic contribution to MT DNA abundance ^14–18^, most of which include a low number of individuals. Therefore, we aimed to leverage data from the UK Biobank, a cohort of more than 500 000 participants aged 40 and above, to uncover non-modifiable as well as life-style factors associated to MT DNA abundance and to further establish a role of inherited genetics in shaping abundance of MT.

## Results

We estimated the amount/abundance of mitochondrial DNA in blood in 295,150 individuals passing quality control (**Supplementary Figure 1** and **Table 1**) from the weighted intensities of genotyping probes mapped to the MT genome (**Supplementary Table S1**). The weighted MT DNA abundance estimate was positively correlated to the mean mitochondrial exome sequencing coverage with a spearman rank correlation coefficient of 0.33 (**Supplementary Figure 1**). We noticed that the individual genotyping batches (each containing around 5,000 individuals) were contributing to the variance of the estimated MT DNA abundance (**Supplementary Figure 2**). In addition, we found a strong effect of genotyping missingness (slope per S.D. and 95% confidence intervals (CI): 0.011 [0.008; 0.015], P<10^−10^) as well as female sex (slope and 95% CI: -0.05 [-0.057; -0.043]; P<10^−43^) on the abundance of mitochondrial DNA in UK Biobank participants. Hence, we present the results separately for women and men, and adjusted our analyses by both the genotyping batch as well as genotyping missingness.

**Table 1.** Baseline characteristics of UK Biobank participants by sex

| Variable | UK Biobank participants |  |  |
| --- | --- | --- | --- |
|  | Females | Males | Both |
| Number of individuals | 158,762 | 136,388 | 295,150 |
| Average age (S.D.) [years] | 56.67 (7.90) | 57.14 (8.09) | 56.89 (7.99) |
| Average body mass index (S.D.) [kg/m<sup>2</sup>] | 27.03 (5.12) | 27.83 (4.21) | 27.40 (4.74) |
| Average packyears smoked (S.D.) | 6.40 (12.89) | 11.05 (18.85) | 8.52 (16.05) |
| Average MT DNA abundance (S.D.)* | 0.02 (0.99) | -0.03 (0.99) | 0.00 (0.99) |
| Median frailty index (IQR) ** | 0.11 (0.01-0.21) | 0.11 (0.01-0.20) | 0.11 (0.01-0.21) |
| Median call rate (range) | 1.00 (0.99-1.00) | 1.00 (0.99-1.00) | 1.00 (0.99-1.00) |

\* Average MT DNA abundance (mL2RMT) estimated from the weighted genotyping probe intensities (L2R), scaled to have a mean of zero and a S.D. of 1.00 in each genotyping plate
\*\* Median fraction of reported health related deficits out of 49 investigated deficits
S.D. = standard deviation; IQR = interquartile range

### Association of mitochondrial DNA abundance with life-style factors and aging

Prior studies implicated a role of age, smoking as well as BMI on Y chromosome loss. Therefore, we first studied how those factors are associated to the MT DNA abundance ^11–13^. Similar to the aforementioned studies, we found increasing age, more packyears as well as elevated BMI to be negatively associated to MT DNA abundance in both men and women (Q-Value<0.05, **Figure 1**).

**Figure 1.**
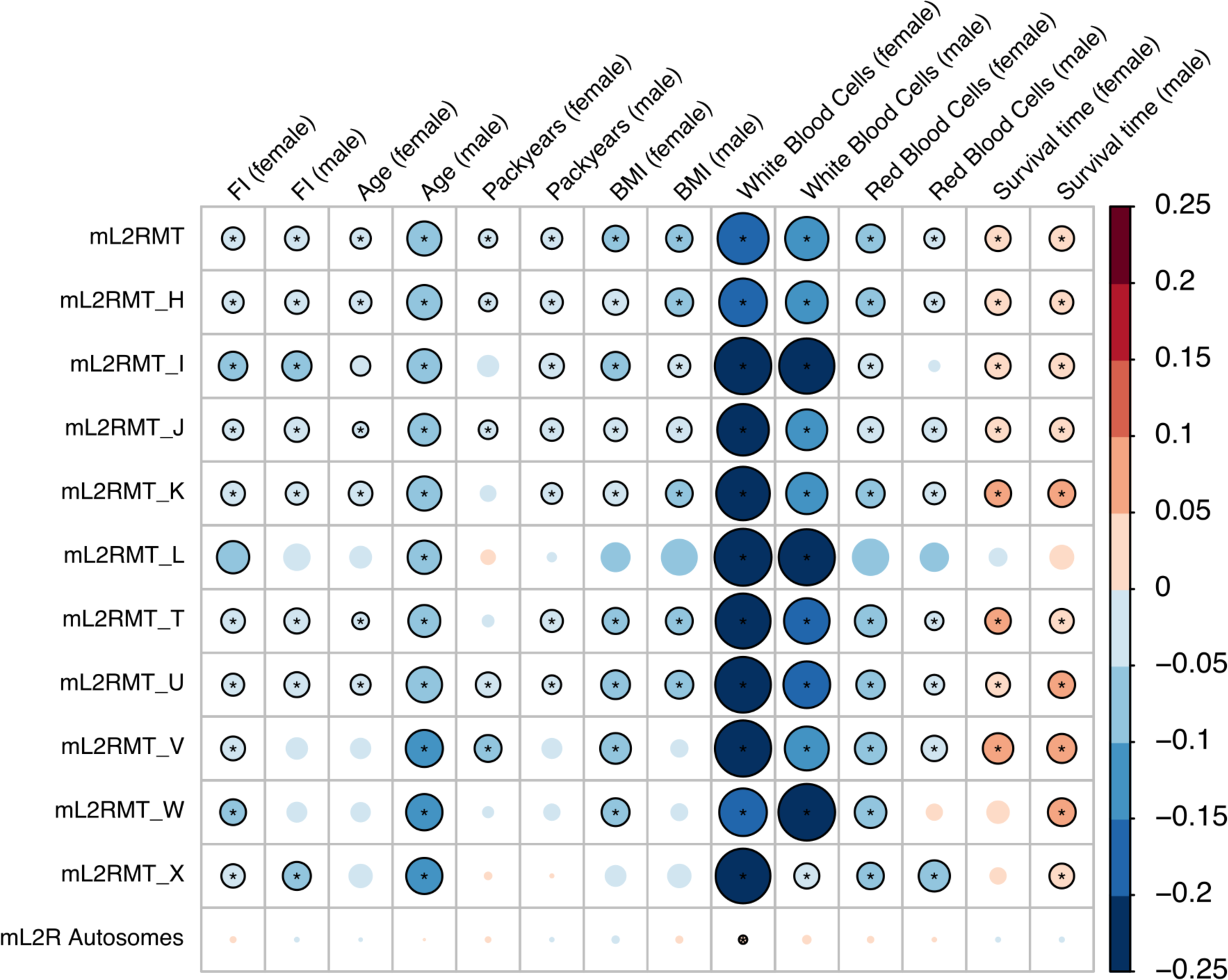
Association between mitochondrial DNA abundance and different covariates. The size and colour (see colour bar) of the circle represent the slope of the association of seven features with mitochondrial (mL2RMT) as well as autosomal DNA abundance (mL2R Autosomes), stratified by sex. The associations were also analysed separately by major MT haplogroup (labelled H-X). Correlations which were statistically significant (Q-Value<0.05) are indicated with an asterisk. Nominally significant associations (P-Value<0.05) are shown as circles with black borders. FI, frailty index; BMI, body mass index; mL2RMT, median log 2 ratio of mitochondrial probes; survival time, time till death or end of follow-up.

In addition, we found that MT DNA abundance is negatively associated to the increased frailty as expressed by the frailty index, an accumulation deficit model summarizing self-reported symptoms and diagnoses (Q-Value<0. 05, **Figure 1**). In line with those results, individuals with higher MT DNA abundance had a statistically significant better/longer survival time after recruitment. Similar results were obtained for the most common haplogroups present in the UKB, indicating that the genetic make-up of the MT itself is not a major factor in determining MT DNA abundance changes due to age or other factors (**Figure 1**).

### Mitochondrial abundance and blood markers

Next, we investigated whether the amount of DNA that is hybridized to the genotyping arrays may be directly related to the number of nucleated cells per volume of blood. Indeed, we found that the white blood cell count was negatively associated to MT DNA abundance (**Figure 1**). However, we also found that increased red blood cell count was associated with reduced MT DNA abundance, indicating that not only the number of nucleated cells seems to play a role in governing MT DNA abundance. This conclusion is supported by the finding that we only observed a weak positive correlation of the aggregated autosomal DNA abundance with white blood cell count, which was estimated from all genotyping probes mapped to autosomes (**Figure 1**). We also investigated individual blood markers and their association with MT DNA abundance and found that the MT DNA abundance was negatively linked to neutrophil, eosinophil, basophil and monocyte percentage and positively linked to lymphocyte percentage and platelet count (**Figure 2A**). In contrast, autosomal DNA abundance was only significantly positively associated with lymphocyte percentage in females.

**Figure 2.**
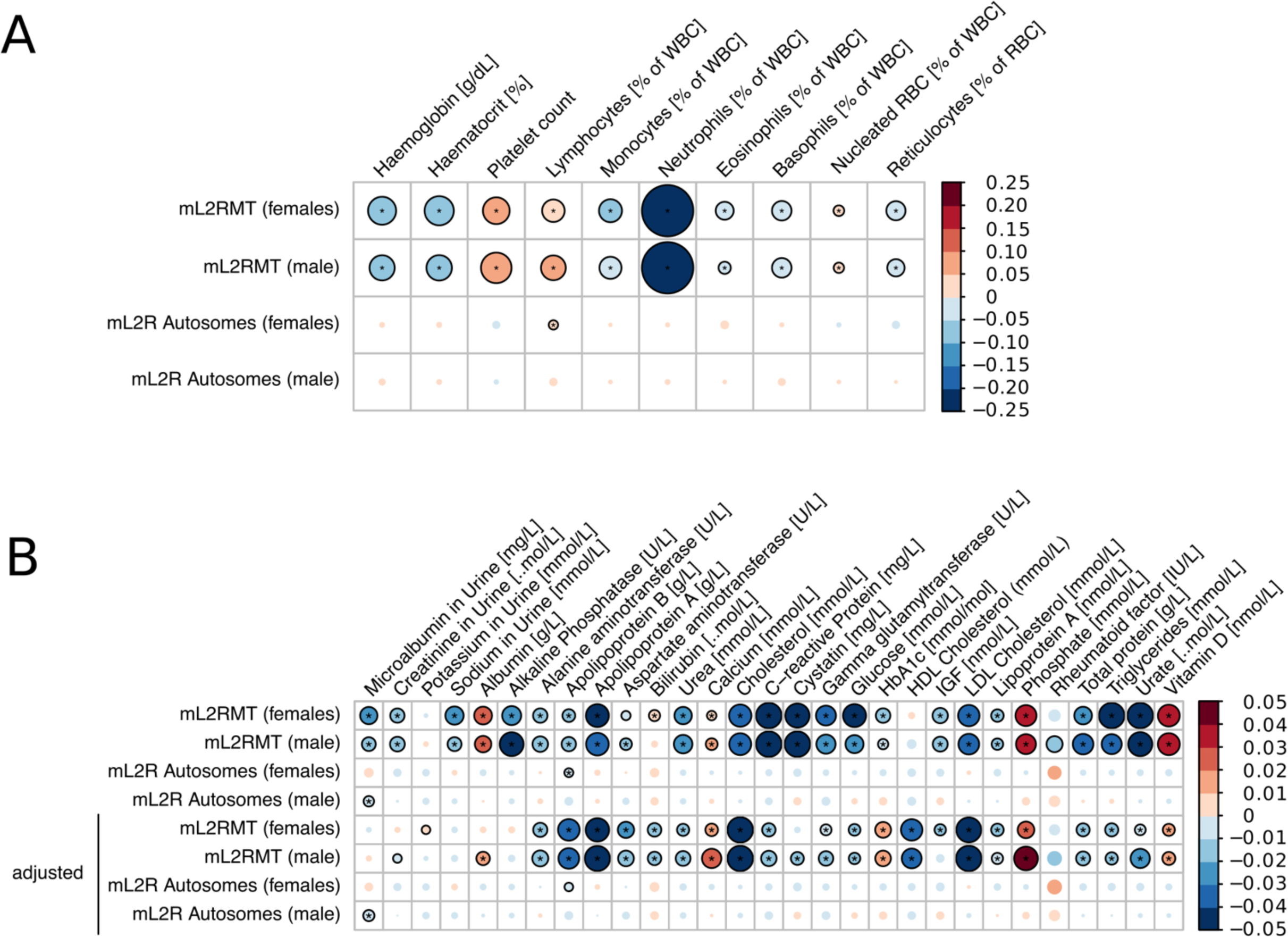
Association between mitochondrial DNA abundance and individual blood markers. The size and colour of the circle represent the slope of the correlation of blood cell counts or other blood measurements (**Panel A**) and biochemistry features (**Panel B**) with mitochondrial (mL2RMT) as well as autosomal DNA abundance (mL2R Autosomes), stratified by sex. In **Panel B**, the association of biochemistry markers with MT DNA abundance were additionally adjusted for the neutrophil and lymphocyte percentage (% of white blood cells) as well as white blood cell count in the last four rows. Correlations which were statistically significant (Q-Value<0.05) are indicated with an asterisk. Nominally significant associations (P-Value<0.05) are shown as circles with black borders. WBC = White blood cells, RBC = red blood cells, mL2RMT, median log 2 ratio of mitochondrial probes, NRBC=Nucleated red blood cells

### Mitochondrial abundance and biochemistry markers

When we investigated the association of 29 biochemistry markers (from urine and blood, **Figure 2B**) and MT DNA abundance, we noticed significant associations with markers related to inflammation (C-reactive protein), kidney function (albumin, cystatin c, urea and urate as well as sodium in urine), liver function (alkaline phosphatase, alanine aminotransferase), cholesterol metabolism (LDL cholesterol, total cholesterol, triglycerides, apolipoprotein A) as well as ion homeostasis (calcium and phosphate), vitamin D levels and glucose metabolisms (IGF, glucose, HbA1c). Since the immune system and thus the number of immune cells may well influence the observed associations, we also present the analyses adjusted for the neutrophil and lymphocyte percentage as well as white blood cell count measured in each participant (**Figure 2B**). After adjustment, most of the associated markers remained statistically significantly associated with MT DNA abundance, albeit with lower effect sizes. Of note, markers related to cholesterol metabolisms such as LDL and HDL cholesterol, apolipoprotein A and B as well as total cholesterol had even greater effect sizes after adjustment for immune cell count. In contrast, we found few associations of biochemistry markers with autosomal DNA abundance in our analyses. In particular, only microalbumin in urine was found to be negatively associated to autosomal DNA abundance in men (Q-Value<0.05).

### Genetic dissection of mitochondrial abundance

To uncover genetic variants associated with mitochondrial abundance, we conducted a genome-wide association study (GWAS) in all individuals passing quality control. In total, we assessed the association of 3,505,788 variants with MT DNA abundance and found 50 regions in the genome with genome-wide significant variants (P-Value<5×10^−08^, **Figure 3**). In those regions, a total of 66 independent signals were detected at genome-wide significance (**Supplementary Table S2**), implicating multiple genes to be involved in MT DNA abundance (**Figure 3**). Importantly, by positional mapping of likely functional variants, we identified genes related to the immune system (*CXCL6, MEF2C, ITPR3, UBE2D1, C7orf73, STIM1, PNP, CRK* and *SIRPB1*), cancer and cell cycle (*TERT, BAK1, CDK6, CDK10, SUFU, FANCI, MDFIC, JMJD1C, USP7, BIK*) and mitochondrial function (*MFN2, TFAM, DGUOK, USP30, CREB5, POLG*) to be potentially involved in governing MT DNA abundance in blood (**Figure 3**).

**Figure 3.**
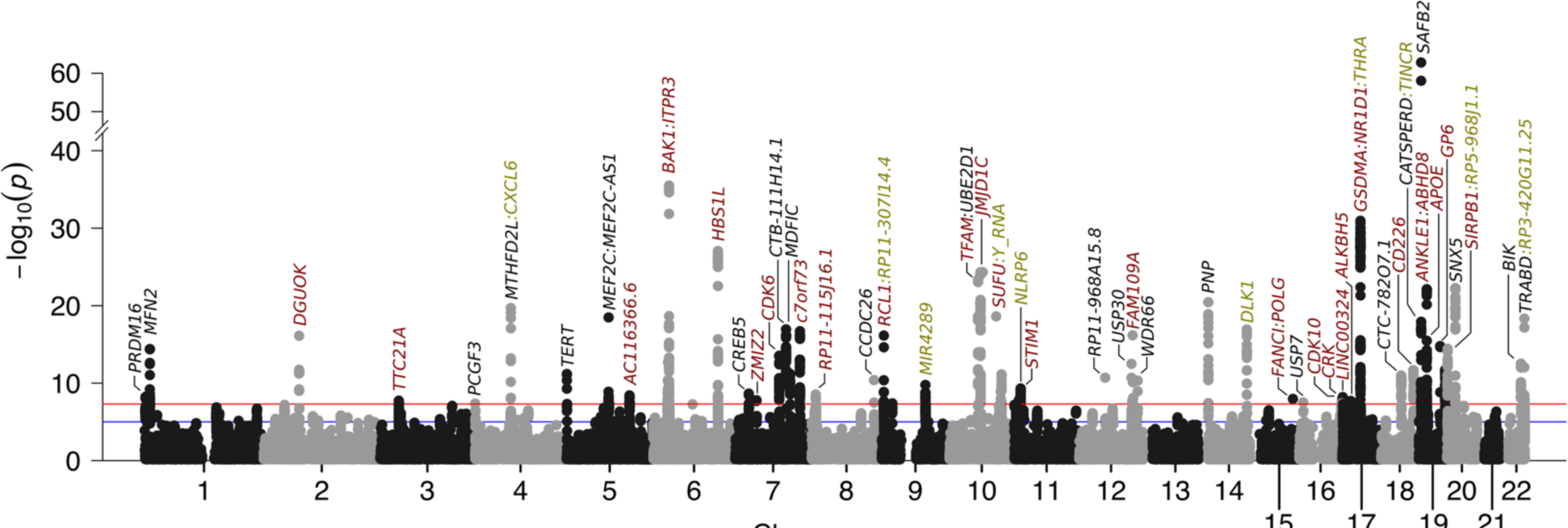
Summary of the genome-wide association study of MT DNA abundance. Manhattan plot of 3,505,788 variants with minor allele frequency greater 5%. Genes influenced by associated variants are highlighted in different colours: genes in red harbour exonic variants, genes which have associated variants in the intronic region are shown in black. In case no intronic or exonic variants are mapped to genes in a region, we highlighted the closest gene in green. The red line denotes genome-wide significance (P<5.00×10^−08^) while the blue line denotes suggestive evidence for association (P<1.00×10^−05^). The estimated genomic inflation factor λ was 1.147.

Next, we used fastBAT to investigate the burden of MT DNA abundance associated variants within the gene bodies of all protein coding genes. In total, 227 genes carrying a statistically significant burden were identified (P-Value<2×10^−06^, **Supplementary Table S3**). We then investigated a potential overrepresentation of pathways within those genes with WebGestaltR. With this approach an overrepresentation of pathways related to immune activation, cell-cell adhesion, haematopoiesis, apoptosis, platelet production as well as mitochondrial biogenesis and plasma lipoprotein assembly was observed (**Supplementary Figure S2** and **Supplementary Table S4**).

Finally, we used mixed effects models to estimate the heritability of MT DNA abundance that can be explained by the genotyped variants. Accounting for age, sex, genotyping missingness, genotyping batch, the first ten genotyping PCs, neutrophil and lymphocyte percentage as well as white blood cell count, we estimated the SNP-heritability to be 8.3% (95% CI: 7.6%-9.0%), strongly implicating a role of inherited genetic variants on MT DNA abundance.

### PheWAS of mitochondrial abundance

In order to estimate the consequence of altered MT DNA abundance on disease risk, we performed a Phenome-wide association study (PheWAS) on ICD10 codes derived from the hospital episode spell data mapped to a total of 1,150 phecodes. We found that higher levels of MT DNA abundance were statistically significantly (Q-Value<0.05) associated with increased risk for (chronic lymphoid) leukaemia, myeloproliferative disease as well as diseases of the spleen (**Figure 4**). In contrast, higher MT DNA abundance resulted in reduced incidence of cardiomegaly, portal hypertension as well as oesophageal bleeding (**Figure 4**).

**Figure 4.**
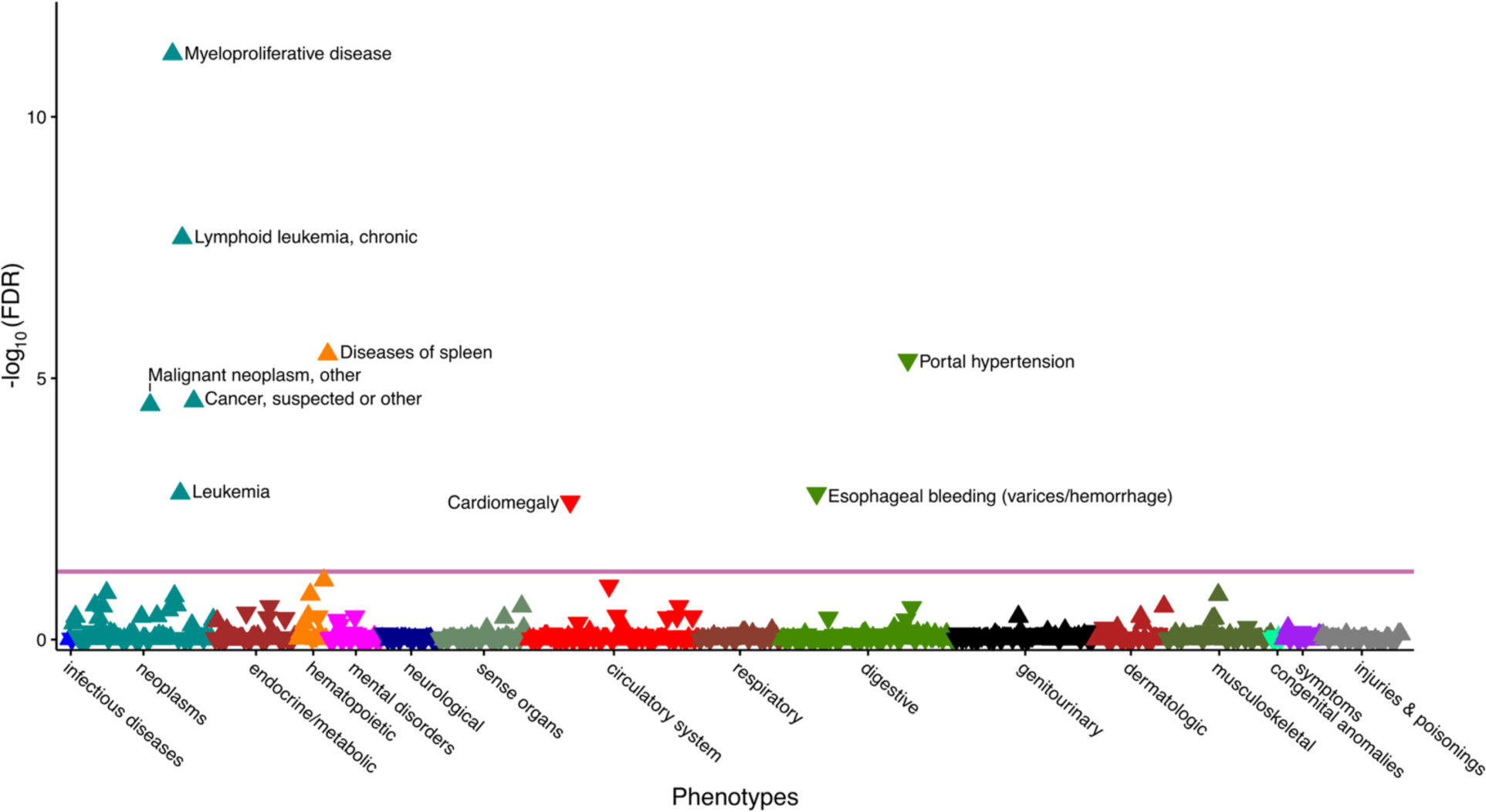
Phenome-wide association study of MT DNA abundance. The consequence of altered MT DNA abundance on 1,150 incident diseases or traits determined from the hospital episode spell data was evaluated in all UK Biobank participants. ICD10 diagnoses were mapped to phecodes and the association between discretized MT DNA abundance (in tertiles) and the phecodes was estimated with logistic regression, adjusted for age at baseline, genotyping batch, total autosomal DNA abundance, fraction of neutrophils, fraction of lymphocytes, white blood cell count, sex, BMI and smoking at baseline as well as the first two principal components of ancestry. Only phecodes with more than 250 cases were considered. Statistically significant associations (Q-Value<0.05) are shown above the red line. Triangles facing up indicate increased risk while triangle pointing down indicate reduced risk to due elevated MT DNA abundance.

## Discussion

In this study, we have explored the epidemiological and genetic basis of MT DNA abundance in nucleated blood cells. We found that age as well as multiple lifestyle and health-related factors were associated with the amount of MT DNA in blood. In particular, MT DNA abundance seems to be mostly influenced by cells involved in the innate immune system as well as cholesterol metabolism and ion homeostasis. In addition, our genetic dissection of MT DNA abundance revealed 50 independent GWAS signals as well as a heritability of around 8%. Finally, we showed that elevated MT DNA abundance is associated with risk for incident leukaemia, hypertensive disorders and oesophageal bleeding.

Due to the low number of variants and thus probes measuring the abundance of the MT genome, we expected that the MT DNA abundance estimates from chip arrays to be noisy. Indeed, we found that our initial MT DNA abundance values were not positively correlated to the coverage of the MT genome by exome sequencing. Since prior studies suggested that exome coverage provides a more robust estimate of MT DNA abundance and also has higher power to detect differences ^19^, we therefore weighted the L2R values of individual array probes to more closely resemble the MT DNA abundance estimates from exome sequencing. With this approach, we found that both estimates are moderately positively correlated (R=0.33) and that the influence of genotype missingness on the weighted MT DNA abundance estimate was reduced more than 10-fold (slope per S.D. and 95%CI of genotype missingness correlated to the mL2RMT estimate from MoChA: 0.146 [0.143; 0.150], P<10^−250^).

Although the effect of genotype missingness on MT abundance estimation is strongly reduced with our approach, it could still potentially confound our analyses. As such, we adjusted our analyses for the call rate to account for those differences. Therefore, it is unlikely that the genotyping missingness after adjustment would strongly confound most of our analyses, particularly since genotyping missingness itself was not associated with any of the investigated markers in this study (Q-values > 0.05, data not shown). In order to estimate the SNP-heritability of MT DNA abundance, the BOLT-LMM algorithm computes the relationship between individuals from all genotyped variants passing QC. Unless genotyping missingness is more strongly pronounced in more related individuals, the estimation of relatedness should not be majorly confounded, particularly after adjusting for the actual genotyping missingness. Our results thus strongly implicate that the amount of mitochondrial DNA in nucleated blood cells is heritable. Further genetic studies accounting for missingness, DNA quality as well as immune cell count are warranted to uncover the precise genetic changes responsible for alternations in MT DNA abundance.

Importantly, our results indicate that MT DNA abundance is negatively associated with increased age, higher frailty, increased smoking behaviour as well as increased BMI. We hypothesized that MT copy number may be diminished due to lower rates of replication with increasing age ^20, 21^, due to oxidative stress induced by smoking ^22^, by low-grade inflammation and metabolic stress due higher BMI ^20^ or due to prior health related conditions (as expressed by the frailty index) ^23, 24^. Since individuals with a healthier lifestyle tend to have higher MT DNA abundance, we also observed a reduced incidence of hypertensive disorders such as portal hypertension and cardiomegaly as well as lower mortality in those individuals. In our GWAS, genes related to blood pressure (*NLRP6, CREB5* and *APOE*) were found in regions associated with MT DNA abundance, providing a potential explanation between increased MT DNA abundance and reduced risk for hypertensive diseases. Similarly, pathways and genes related to coagulation, platelet production and count were implicated by our GWAS, thus pointing towards genes that are involved in both MT DNA abundance and risk for oesophageal bleeding. Interestingly, recent reports indicated that higher MT abundance is protective against oesophageal cancer ^25^, which may indicate that the oesophageal tissue is particularly vulnerable to changes in mitochondrial DNA abundance.

To further reinforce the notion that MT DNA abundance is heritable, we conducted a genome-wide association study on over 3 million common variants. We chose to focus on common variants, since they are less likely to suffer from missing genotypes and imputation of missing alleles is more accurate for those variants as well. Our GWAS implicated several genes to be involved in governing MT DNA abundance, particularly those related to immunity, cancer, cell cycle and mitochondrial function. This is in line with reports that the loss of Y chromosome (mLOY) is related to similar pathways ^26^, although none of the genes implicated in this study overlap with the genes associated with mLOY. Furthermore, we were able to identify variants significantly associated with MT DNA abundance within a region on chromosome 17 (association signal near GSDMA, i.e. genomic locus 40 in **Supplementary Table S2**), which was previously shown to be involved in MT DNA abundance (estimated from quantitative PCR ^15^). This regions was also described to be involved in blood cell counts, particularly neutrophils ^27–29^. Therefore, our approach does seem to accurately capture MT DNA abundance and provides evidence for additional regions in the genome and thus genes involved in MT DNA abundance.

We observed a strong negative association between MT DNA abundance and the number of neutrophils and a positive correlation with lymphocytes percentage. Since neutrophils are the most abundant leukocyte and thus the most abundant cell carrying DNA, it may seem obvious that the amount of DNA hybridized to the array should be positively correlated. However, our results rather point to other conclusions. First, we did not observe an association of autosomal DNA abundance with neutrophil count, indicating that the total amount of DNA that was hybridized does not play a major role, potentially due to careful harmonization of DNA concentration used in the genotyping process. Second, we observed differential correlation for MT DNA abundance with myeloblast-derived and lymphoid immune cells. Since both immune cell lineages carry mitochondria, they should similarly contribute to increased MT DNA abundance in case the measured MT DNA abundance is solely dependent on DNA content in blood. Therefore, we conclude that other processes must be responsible for the observed association between increased MT DNA abundance and reduced (innate) immune cell count.

In our PheWAS analysis, we found increased MT DNA abundance to be associated with increased risk of leukaemia, which is in agreement to prior findings which found higher MT DNA abundance to cause increased leukaemia risk ^30, 31^. Since an increased number of circulating lymphocytes is a hallmark of lymphoid leukaemia ^32^ and we found a positive association of MT DNA abundance and lymphocyte count, this association would seem obvious. However, we adjusted our PheWAS by neutrophil and lymphocyte percentage as well as white blood cell count, thus the observed effect should be largely independent of immune cell abundance, implicating that other mitochondrial mechanisms independent of white blood cell count are responsible for the increased risk. Importantly, in our GWAS, which was also adjusted for white blood cell counts, we found association signals near genes involved in leukaemia and lymphoma development (*CRK, BAK1, CDK6, MDFIC, BIK*) as well as a significant enrichment of pathways related to immune cell proliferation. Thus, further functional studies are necessary to identify the underlying mechanism and responsible (immune) cell population that links the function of those genes to MT DNA abundance and leukaemia.

Multiple blood and urine bound biochemistry markers related to cholesterol and triglyceride metabolism as well as ion homeostasis and vitamin D levels were found to be associated with MT DNA abundance independent of neutrophil and lymphocyte percentage or white blood cell count. This is in line with the proposed function of mitochondria in cells since they are instrumental in the metabolism of cholesterol and other lipids as well as in ion storage, particularly of phosphate and calcium.

A recent study presented evidence that cell-free and respiratory competent mitochondria are present in blood and may have a role in cell-cell communication ^33^. Although the majority of DNA used for the genotyping experiments should be derived from nucleated cells, our current approach can, however, not distinguish between cellular and cell-free mitochondria. It is therefore possible that some of the observed associations are additionally driven by extracellular mitochondria, which warrants further investigations.

## Conclusion

We show that the amount of MT DNA in nucleated blood cells is heritable and associated with different modifiable and non-modifiable markers related to aging, health and lifestyle as well as immunity, cholesterol metabolism and ion homeostasis. Our findings suggest that estimating MT DNA abundance from weighted intensities on genotyping arrays is a promising tool to gain novel insights into disease relevant mechanisms as well as to study aging.

## Methods

### UK Biobank study population

UK Biobank (UKB) is a multi-centre cohort with 502,621 participants recruited in England, Scotland and Wales. Genotype data was available for 488,279 participants ^34^. In addition to the quality control steps by the UKB, we had to apply additional exclusion criteria to the UK Biobank participants (**Supplementary Figure S1**). As such, we further excluded individuals with documented and inferred sex chromosome abnormalities and those that had a low call rate (less than 99.0%) or excessive heterozygosity as determined by the UKB. Finally, we restricted the analyses to British Caucasians (determined by the genotype principal components) as well as unrelated individuals, resulting in a total analytical dataset of 295,150 samples (**Table 1**). Further quality control measures based on the genetic dosage/intensity estimation are indicated below. The current study was conducted as part of the registered project 22224.

### Variables evaluated

The age and BMI of the individuals was recorded at the time of blood draw and smoking status were ascertained at baseline interview via questionnaire. The frailty index (FI) consisting of 49 health-related items, was computed as recently described ^35^ for all participants from questionnaire data. Briefly, the Rockwood accumulation deficit model was applied summarizing the number of self-reported diseases, symptoms and signs as well as overall health assessments in each individual. The FI ranges from zero to one and represents the fraction of actual number of health deficits reported by an individual divided by the total number of potential health deficits considered (i.e. queried in the questionnaire). Increased frailty of an individual represents a major risk factor for future diseases as well as mortality. Recently, the UKB measured a panel of 29 biomarkers in urine and blood and the data were released to the public after extensive quality control. In addition, platelet, red and white blood cell counts were measured as an absolute number per unit volume and the abundance of individual blood cell classes such as lymphocytes, monocytes, neutrophils, eosinophils, basophils were recorded in percent (%) of white blood cells. Calibration and quality control were performed by the UK Biobank investigators. All-cause mortality and thus survival time after recruitment was ascertained from the date and cause of death reported up until 2018-02-11 for all deceased UK Biobank participants (mean follow-up time: 8.90 years).

### Computation of mitochondrial abundance in UK Biobank participants

In the UKB participants, we determined somatic mitochondrial DNA abundance from the intensities of genotyping probes on the MT chromosome. On Affymetrix arrays, the relative amount of DNA hybridized to the array at each probe is expressed as the log2 ratio (L2R), which represents the log2 transformed ratio of the observed genotyping probe intensity divided by the intensity at the same probe observed in a set of reference samples.

Initially, we computed the median L2R values across all 265 variants on the MT chromosome with MoChA ^36^. To compare those estimates to another method of MT DNA abundance determination, we also computed the average coverage of the MT genome as estimated with mosdepth ^37^ in around 50,000 individuals with exome sequencing data in the UK Biobank ^38^. However, we noticed that the average MT coverage, normalized to the total coverage was negatively correlated to the MT DNA abundance values acquired with MoChA (R=-0.07). This finding indicated that several poorly performing probes were confounding the abundance estimation. We therefore fit a multivariate linear regression model to select probes that are statistically significantly predicting the normalized MT coverage from exome sequencing data (**Supplementary Table S1**). We then multiplied the respective L2R value of each probe by the weight in **Supplementary Table S1** and computed the median L2R value across those weighted L2R values (mL2RMT), resulting in a single MT DNA abundance estimate for each individual. The distribution was rescaled to have a mean of zero and a standard deviation (S.D.) of one within each genotyping plate consisting of 96 samples. To further analyse potentially haplogroup specific effects, haplogroups in the UKB participants were estimated with *haplogrep2* (version 2.1.25) ^39^ from all MT variants genotyped on the chip with standard settings. We only analysed the top-level haplogroups that were present in more than 1,000 individuals due to power considerations.

### Additional quality control based on intensity and whole exome sequencing data in the UKB

Similar to the quality control steps suggested for Illumina genotyping arrays, for each individual, we used MoChA to compute the standard deviation of the L2R values (SDL2R) of all autosomal variants and only included individuals with SDL2R of less than 0.36 (i.e. less than two S.Ds from the mean SDL2R of all samples), effectively excluding samples with excessive variation in their genotyping intensities. Furthermore, we excluded individuals with an b-allele frequency phase concordance across phased heterozygous sites greater than 0.52 ^40^. In addition, we also computed the average autosomal DNA abundance from all autosomal L2R values (mL2Rauto) with MoChA. Adjusting for this variable allows us to account for differences in genotyping quality and differences in DNA hybridization to the genotyping chip. Furthermore, we compared the association results from the MT DNA abundance to the overall (autosomal) DNA abundance to identify MT specific effects.

### Statistical analyses

We coded the continuous MT DNA abundance (mL2RMT) as the outcome and used linear regression in R (function *lm* as implemented in base R) to evaluate its correlation with exposure variables. All analyses were adjusted for age at baseline/blood draw, sex (determined from the genotyping data), the first two genotype principal components (PCs), the number of missing genotypes as well as genotyping batch. We also performed some association testing by stratifying patients according to their MT haplogroup. Those analyses were additionally adjusted by the quality (i.e. confidence) of the respective haplotype computation as reported by *haplogrep2*. Unless otherweise mentioned, to account for multiple testing (where appropriate), we controlled the false-discovery rate to be less than 5% and thus report results that have a Q-Value of less than 0.05.

### Genome-wide association study of mitochondrial abundance

The genome-wide association study (GWAS) of mitochondrial DNA abundance was computed with plink2 ^41^. We restricted the analyses to variants with a minor allele frequency greater than 5%, with less than 2% missing data, a MACH R^2^ imputation quality greater than 0.6 as well as not deviating significantly (P<10^−05^) from Hardy-Weinberg equilibrium. The association between MT DNA abundance and the additive genotypes in the genome-wide association study was adjusted for by the first 10 genotype PCs, age at baseline, sex, genotyping batch, genotyping missingness/call rate as well as neutrophil percentage, lymphocyte percentage and white blood cell count. The genomic inflation factor λ was calculated with the *GenABEL* library version 1.8-0 implemented in R. We considered variants associated with MT DNA abundance with a P-value lower than 5×10^−08^ to be genome-wide significantly correlated to MT DNA abundance. Functional mapping of associated variants to genes within genome-wide significant loci was done with FUMA ^42^. Independent signals within each locus were defined as additional association signals at genome-wide significance and with a correlation (R^2^) of less than 0.1 to the variant with the lowest P-Value. In each locus we considered the variant (as well as correlated variants at R^2^ > 0.6) within gene bodies with the highest CADD ^43^ score to be the most likely functional variant and also recorded the affected gene (i.e. labelled each locus according to the gene name). Similarly, for variants outside of the gene body, we extracted the variant with the lowest RegulomeDB ^44^ score as well as the closest gene to that variant.

We used fastBAT implemented in the Complex-Traits Genetics Virtual Lab ^45^ using the GWAS summary statistics to assess the genetic burden of variants associated with MT DNA abundance within the gene body of all genes. The statistical significance threshold was set at P-Value<2.0×10^−6^ which is the Bonferroni correction threshold for 20,000 protein-coding genes. Genes with a statistically significant burden were analysed in a pathway overrepresentation analysis (ORA) with Webgestalt as implemented in WebGestaltR ^46^ in R. Pathways for the ORA were retrieved from MSigDB^47^ version 7.1 and we report pathways with a size of between 25 and 1,000 genes and with a Q-Value of less than 0.05.

### Heritability estimation of mitochondrial DNA abundance

The phenotypic variance explained by additive contributions from SNPs (SNP heritability) of MT DNA abundance was estimated with the genomic restricted maximum likelihood estimation method implemented in the genome-wide complex trait analysis in BOLT-LMM ^48^, with standard settings and adjusted for the same covariates as described for the genome-wide association analysis.

### PheWAS analysis

Diseases were extracted from the hospital episode spell data of all individuals passing QC and the ICD10 codes were mapped to phecodes ^49^ using the *createPhewasTable* function from the *PheWAS* package ^50^ implemented in R. We only considered incident diseases, i.e. diseases that occurred at least once after blood draw/recruitment. We also excluded all patients with prevalent diseases from the respective incident data set so only individuals without pre-existing conditions were evaluated within each respective phecode or phecode grouping. The PheWAS was performed with the *phewas* function and we only considered phecodes which occurred in more than 250 cases. In order to reduce the impact of extreme mL2R values and to enable comparison with prior studies ^30, 31^, we discretized the variable according to tertiles and used this variable as the exposure in the PheWAS. The analyses were adjusted for the same covariates as the genome-wide association study. Results were plotted with the *phewasManhattan* function.

## Data availability

The data were exclusively retrieved from the UK Biobank and can be accessed upon request from the UK Biobank. The mitochondrial DNA abundance as computed in this manuscript will be reported back to the UK Biobank upon publication. The scripts to compute the weights and the weighted MT DNA abundance in the UKB dataset will be published at https://github.com/GrassmannLab/MT_UKB.

## Acknowledgement

This research has been conducted using the UK Biobank Resource under Application Number 22224.

## Author Contributions

**Conception and design:** Felix Grassmann, Sara Hägg

**Financial support:** Kamila Czene, Sara Hägg

**Collection and assembly of data:** Felix Grassmann, Juulia Jylhävä, Sara Hägg

**Data analysis and interpretation:** Felix Grassmann, Juulia Jylhävä, Yunzhang Wang, Sara Hägg

**Manuscript writing:** Felix Grassmann, Kamila Czene, Sara Hägg, Juulia Jylhävä

**Final approval of manuscript:** All authors

---

**Supplementary Figure S1.**
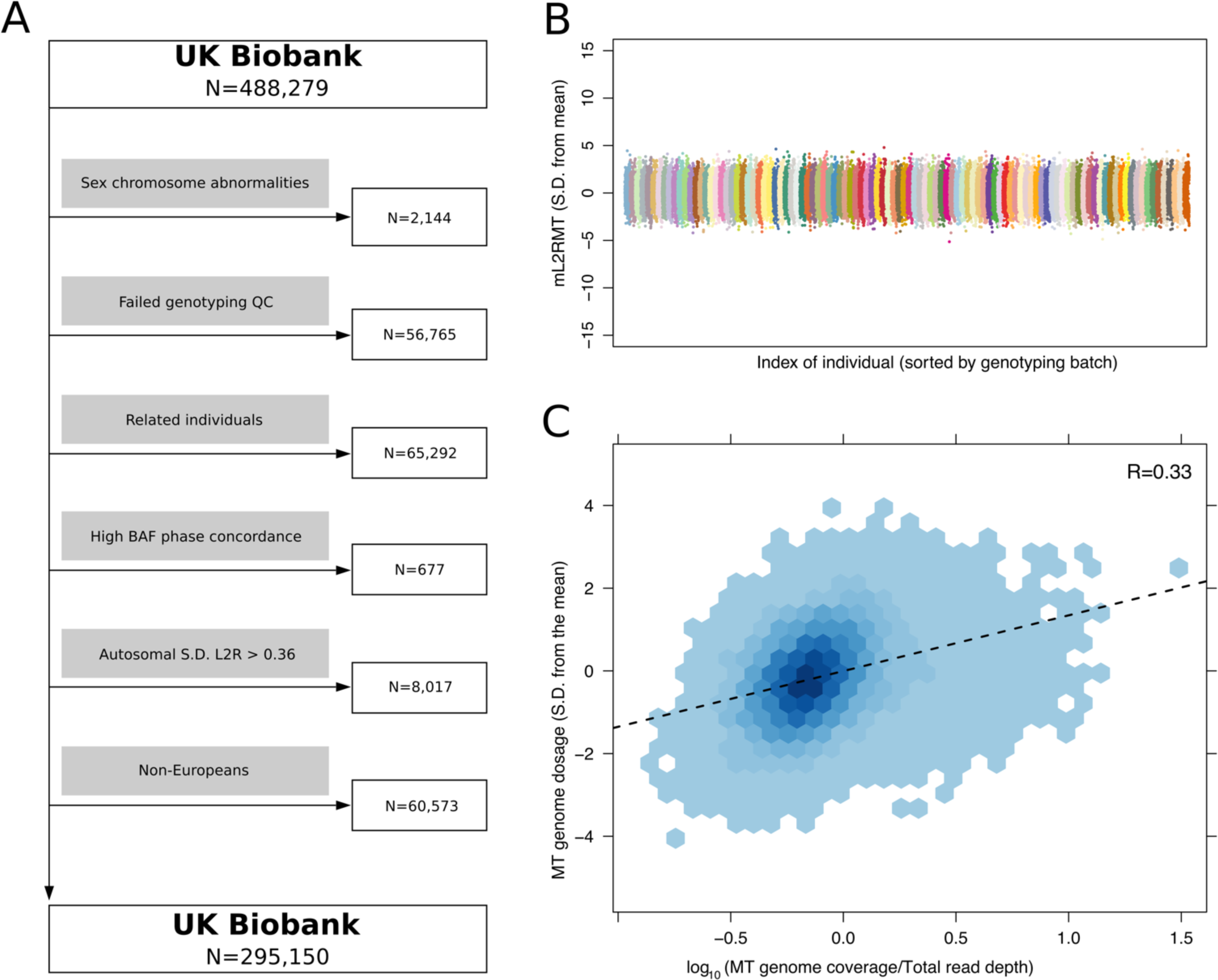
Exclusion criteria and quality control in the UK Biobank. **A.** Exclusion criteria in the UK Biobank according to established and intensity/dosage specific quality control criteria. **B.** Distribution of MT DNA abundance across the whole cohort in standard deviations (S.D.) from the mean, coloured by genotyping batch. **C.** Correlation between MT DNA abundance computed from the weighted genotyping chip intensities and the average coverage of the MT genome (normalized by the total read depth of each individual) in around 50,000 individuals with available exome sequencing data ^38^.

**Supplementary Figure S2.**
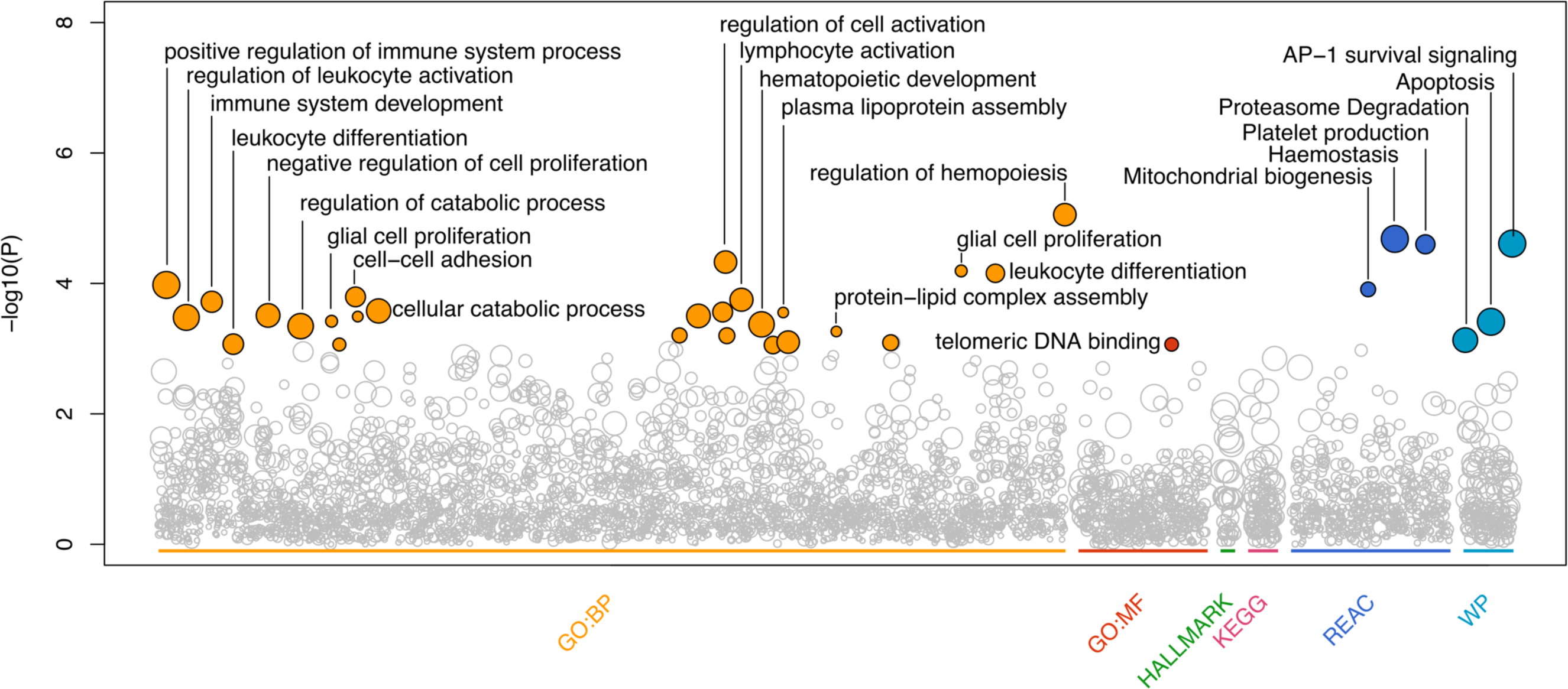
Pathway overrepresentation of genes with a significant burden of common variants associated with MT DNA abundance. A pathway enrichment analyses was performed with WebGestaltR to estimate the overrepresentation of genes with a significant burden due to common MT DNA abundance associated variants. Pathways statistically significantly enriched (Q-Value<0.05) are highlighted with solid circles, those below statistical significance are indicated with a transparent grey circle. Selected descriptions for statistically significant pathways are shown.

**Supplementary Table S1.**
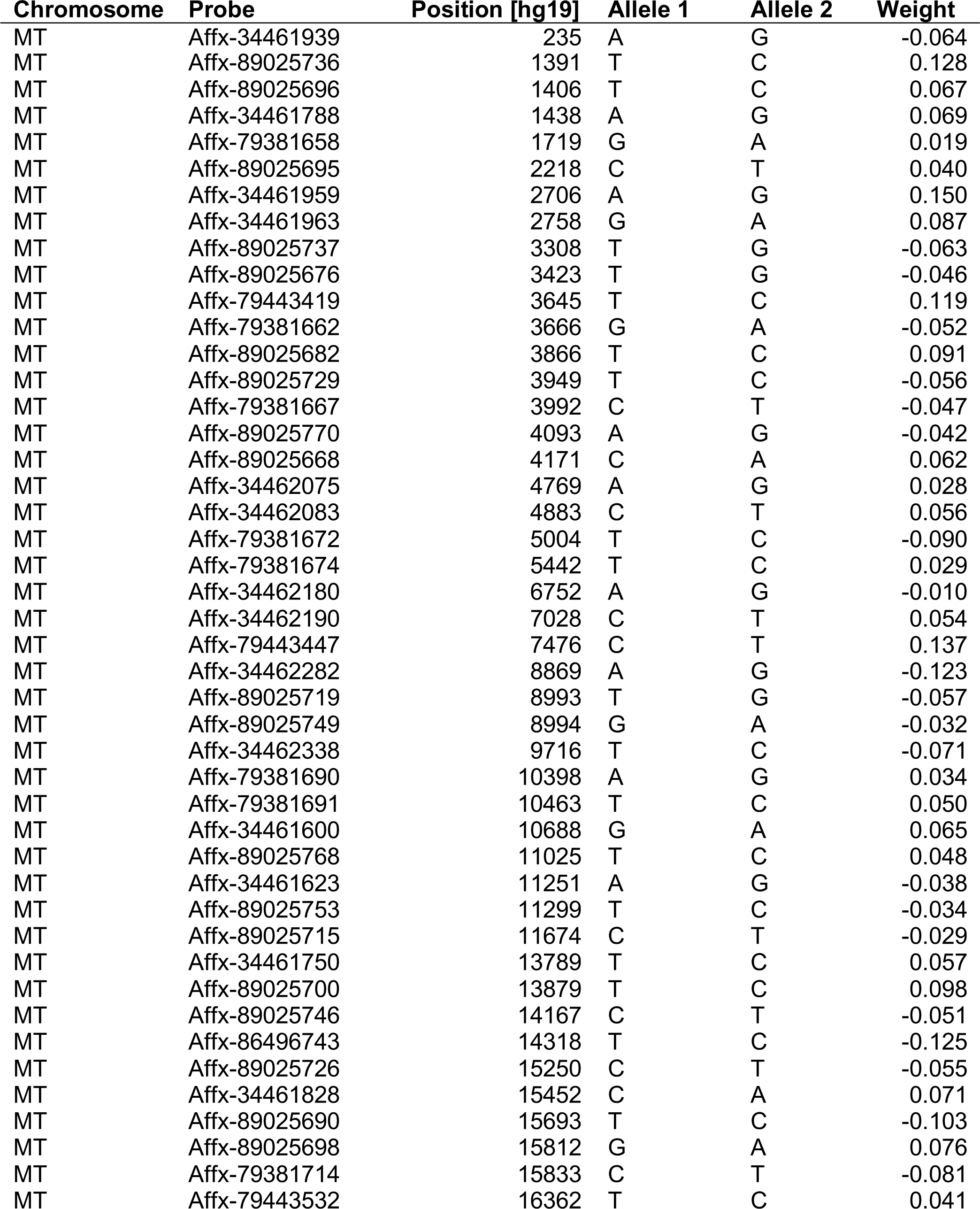
Weights used to compute the MT DNA abundance from genotyping probe intensities (L2R)

| Chromosome | Probe | Position [hg19] | Allele 1 | Allele 2 | Weight |
| --- | --- | --- | --- | --- | --- |
| MT | Affx-34461939 | 235 | A | G | -0.064 |
| MT | Affx-89025736 | 1391 | T | C | 0.128 |
| MT | Affx-89025696 | 1406 | T | C | 0.067 |
| MT | Affx-34461788 | 1438 | A | G | 0.069 |
| MT | Affx-79381658 | 1719 | G | A | 0.019 |
| MT | Affx-89025695 | 2218 | C | T | 0.040 |
| MT | Affx-34461959 | 2706 | A | G | 0.150 |
| MT | Affx-34461963 | 2758 | G | A | 0.087 |
| MT | Affx-89025737 | 3308 | T | G | -0.063 |
| MT | Affx-89025676 | 3423 | T | G | -0.046 |
| MT | Affx-79443419 | 3645 | T | C | 0.119 |
| MT | Affx-79381662 | 3666 | G | A | -0.052 |
| MT | Affx-89025682 | 3866 | T | C | 0.091 |
| MT | Affx-89025729 | 3949 | T | C | -0.056 |
| MT | Affx-79381667 | 3992 | C | T | -0.047 |
| MT | Affx-89025770 | 4093 | A | G | -0.042 |
| MT | Affx-89025668 | 4171 | C | A | 0.062 |
| MT | Affx-34462075 | 4769 | A | G | 0.028 |
| MT | Affx-34462083 | 4883 | C | T | 0.056 |
| MT | Affx-79381672 | 5004 | T | C | -0.090 |
| MT | Affx-79381674 | 5442 | T | C | 0.029 |
| MT | Affx-34462180 | 6752 | A | G | -0.010 |
| MT | Affx-34462190 | 7028 | C | T | 0.054 |
| MT | Affx-79443447 | 7476 | C | T | 0.137 |
| MT | Affx-34462282 | 8869 | A | G | -0.123 |
| MT | Affx-89025719 | 8993 | T | G | -0.057 |
| MT | Affx-89025749 | 8994 | G | A | -0.032 |
| MT | Affx-34462338 | 9716 | T | C | -0.071 |
| MT | Affx-79381690 | 10398 | A | G | 0.034 |
| MT | Affx-79381691 | 10463 | T | C | 0.050 |
| MT | Affx-34461600 | 10688 | G | A | 0.065 |
| MT | Affx-89025768 | 11025 | T | C | 0.048 |
| MT | Affx-34461623 | 11251 | A | G | -0.038 |
| MT | Affx-89025753 | 11299 | T | C | -0.034 |
| MT | Affx-89025715 | 11674 | C | T | -0.029 |
| MT | Affx-34461750 | 13789 | T | C | 0.057 |
| MT | Affx-89025700 | 13879 | T | C | 0.098 |
| MT | Affx-89025746 | 14167 | C | T | -0.051 |
| MT | Affx-86496743 | 14318 | T | C | -0.125 |
| MT | Affx-89025726 | 15250 | C | T | -0.055 |
| MT | Affx-34461828 | 15452 | C | A | 0.071 |
| MT | Affx-89025690 | 15693 | T | C | -0.103 |
| MT | Affx-89025698 | 15812 | G | A | 0.076 |
| MT | Affx-79381714 | 15833 | C | T | -0.081 |
| MT | Affx-79443532 | 16362 | T | C | 0.041 |

**Supplementary Table S2. Genome-wide significant loci associated with MT abundance**

**Supplementary Table S3. Genes with a significant burden of common variants**

**Supplementary Table S4. Pathway overrepresentation analysis (ORA) results**

